# StrainFacts accurately quantifies both endogenous and live biotherapeutic product strain abundances in simulated and clinical vaginal microbiota samples

**DOI:** 10.1101/2025.08.15.670563

**Authors:** Johnathan Shih, Seth M. Bloom, Jiawu Xu, Caroline M. Mitchell, Joseph Elsherbini, Douglas S. Kwon

## Abstract

Live biotherapeutic products (LBPs) deliver microbial strains to modulate the host microbiota in order to promote health or treat and prevent disease. Since endogenous strains are already present, accurately evaluating LBP efficacy and mechanism of action requires distinguishing administered from endogenous strains. Although computational tools exist for inferring strains from short-read metagenomic data, few have been rigorously tested in the context of LBP treatment. Here, we assess the ability of StrainFacts, a computational tool for inferring strains from short-read metagenomic data, to estimate strain abundances and genotypes of endogenous and administered strains. We performed a simulation study of a single-strain LBP trial, modeling serial samples across a range of administered strain abundance, co-occurring endogenous strains, and sequencing depths. We found that StrainFacts accurately estimated both LBP and endogenous strain abundances and genotypes within simulated samples. We further validated methods using human vaginal microbiota samples spiked with CTV-05, the *Lactobacillus crispatus* strain contained in the LBP LACTIN-V, which has been shown to reduce recurrent bacterial vaginosis. Our findings demonstrate that StrainFacts can robustly assess LBP and endogenous strain colonization, abundance, and dynamics in simulated and experimental microbiota samples, supporting its utility as an analysis tool for vaginal LBP therapeutic trial data.

## Introduction

Live biotherapeutic products (LBPs) contain one or more defined bacterial strains administered with therapeutic intent to modify microbiota composition and/or promote host mucosal or systemic health. They represent a promising therapeutic approach for diverse mucosal and microbiota-associated diseases (Cohen 2020, Khanna 2021,Vermeire 2024, Hansen 2021). Because LBPs often contain strains of species also naturally present in the host microbiota, evaluating LBP efficacy, colonization, and mechanisms of action requires precise differentiation between administered strains and endogenous strains of the same species. Accurate strain tracing is therefore essential for assessing colonization success, persistence, and ecological dynamics, yet validated tools for this purpose remain limited. Quantitative PCR (qPCR) is frequently used for LBP strain detection, but its reliance on short target sequences inherently limits strain resolution and can generate false positive results when endogenous strains share identical sequences. In addition, qPCR typically cannot track endogenous strains without developing custom assays after sample collection. Shotgun metagenomics offers the ability to discriminate strains by leveraging entire genomes, but existing bioinformatics tools for strain inference from metagenomic data have not been rigorously validated for LBP applications. An ideal tool would both reliably distinguish administered strains from endogenous strains of the same species and accurately quantify their relative abundances to assess strain dynamics.

Here, we validate a strain deconvolution tool, StrainFacts, for strain-level analysis of LBP data and develop a workflow for analyzing strain dynamics in the vaginal microbiome. StrainFacts uses as input the metagenotype of a target species for each sample (Shi 2023). We first performed a study using simulated metagenotypes spiked with a simulated LBP (sLBP). We then assess the ability of StrainFacts to identify and quantify administered and endogenous strains using metagenomic data from human vaginal samples with varying microbiota compositions, spiked with the *Lactobacillus crispatus* strain CTV-05 - the microbial component of the single-strain LBP LACTIN-V (Cohen 2020). Our analysis demonstrates that StrainFacts effectively tracks both administered and endogenous strains in simulated and real-world vaginal microbiome samples.

## Results

### Review of available tools for bacterial strain analysis

StrainFacts is one of few available strain-tracing tools that infers strain relative abundances and genotypes globally across all samples, and it is modern, currently maintained, and runs quickly even on thousands of samples with GPU acceleration. (**Table S1**). Tools like StrainPhlan (Truong 2017) infer presence and absence of a strain but do not provide relative abundances. Other tools that determine the relative abundance of strains often require higher sequencing depth, which may limit applicability to low abundance strains. StrainFacts simultaneously infers strain genotypes and quantifies relative abundances of strains across all samples, making it suitable for tracking both LBP and endogenous strains in large studies.

### StrainFacts accurately identifies and quantifies a simulated LBP strain

To evaluate the performance of StrainFacts on simulated samples, we first assessed its accuracy in inferring the genotype of a simulated live biotherapeutic product (sLBP) strain. We simulated and analyzed 10 unique single-arm LBP treatment cohorts, each including 100 participants with 4 unique samples per participant, each containing 1 to 4 simulated endogenous strains of the LBP species at varying abundances and sequencing depths, plus a single sLBP strain “spiked-in” at varying abundance (**Figure 1A**). In all 10 simulated cohorts, after combining highly similar strains with a Jaccard Similarity of ≥0.95, a single strain with an inferred genotype identical to the sLBP strain’s true genotype (Jaccard Similarity = 1) was identified (**Figure 1B**), demonstrating highly accurate sLBP genotypic inference.

**Figure 1.**
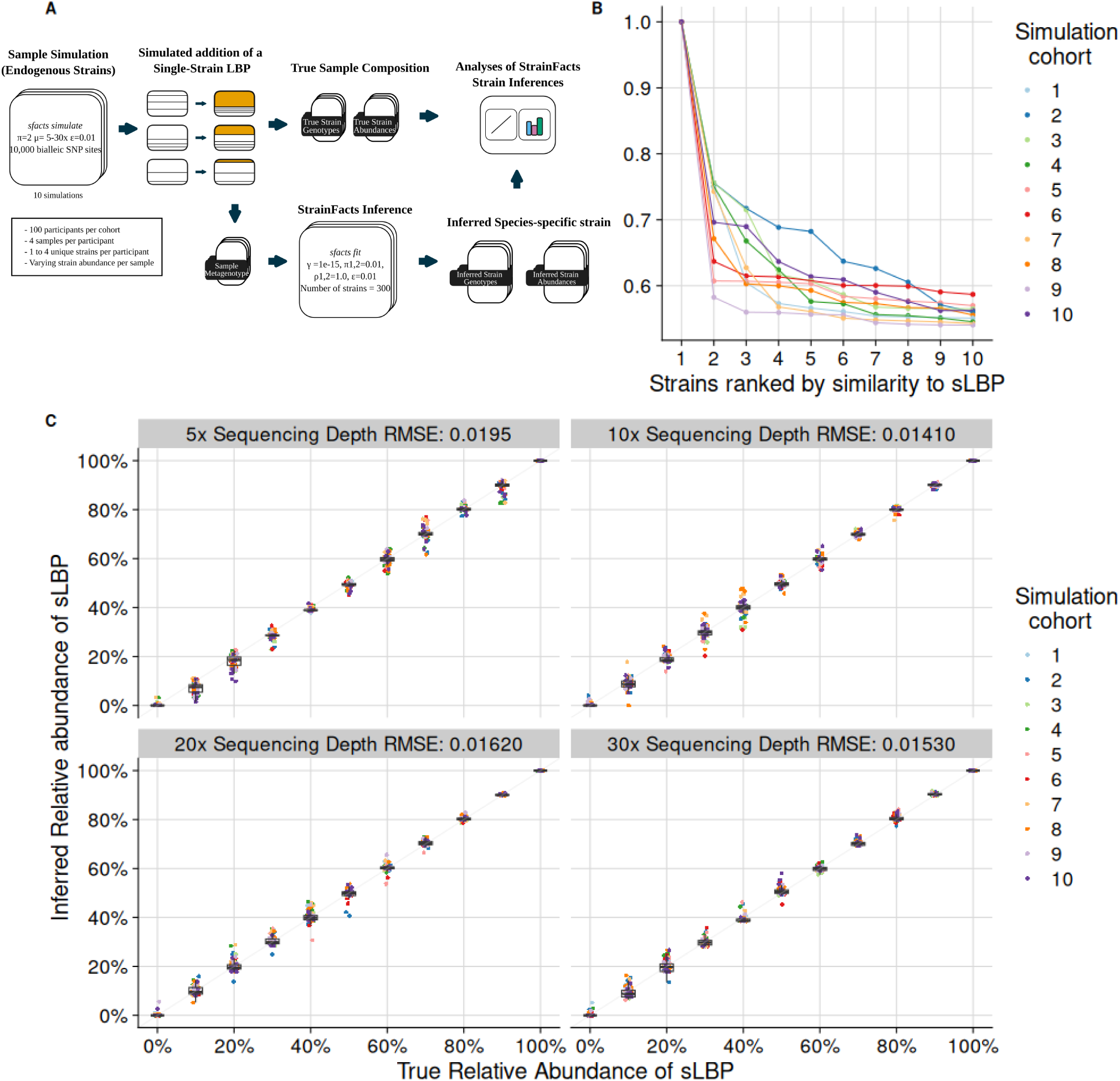
StrainFacts performance in inferring presence, genotype, and abundance of a live biotherapeutic product (LBP) strain in simulated cohorts. A) A schematic of the simulation process for each simulated cohort. B) Jaccard similarity coefficients of the ten strains most similar to the simulated LBP (sLBP) strain’s genotype in each of the 10 simulated cohorts. C) Relative abundances of the inferred sLBP strain in simulated samples across varying sequencing depth. Colors indicate the simulation cohort from which the strain was derived. Root Mean Squared Error (RMSE) for the comparison of true and inferred relative abundances of the simulated LBP strain are calculated for simulated samples at each sequencing depth across all simulated cohorts.

Having confirmed accurate genotypic identification, we next investigated how accurately StrainFacts estimated the sLBP strain’s per-sample abundance. For each sample, the inferred abundance of the sLBP strain was compared to its true abundance, with inference accuracy determined by calculating the Root Mean Square Error (RMSE) for each simulated cohort at each sequencing depth. Across all cohorts, StrainFacts accurately inferred sLBP strain per-sample abundance as demonstrated by low (<0.02) RMSE values at all sequencing depths, with greater depths yielding superior performance (**Figure 1C**). Across all simulations, 4.44% of samples with a true sLBP strain abundance of 0% exhibited inferred sLBP abundance >1% abundance (“false positives”), although the false positives were all estimated at <10% per-sample abundance, while 3.33% of samples with true sLBP abundance of 10% had inferred sLBP abundance <5% abundance (“false negatives”). Based upon these results, we applied a ≥10% inferred abundance threshold to define strain presence in subsequent analyses, thereby reducing the false positive rate.

### StrainFacts accurately infers genotypes and abundances of simulated endogenous strains

In addition to quantifying presence and abundance of the LBP strains, it is also important to characterize endogenous (“native”) strains from the same species to more fully capture LBP treatment effects on microbiota composition and strain dynamics. We investigated StrainFacts’s accuracy in inferring endogenous strains by computing the Jaccard similarity between each true strain’s genotype and each inferred strain’s genotype, and assessing how well they matched.

We examined the effect of varying the Jaccard similarity threshold value to define matches between true and simulated strain genotypes. An optimal threshold would match each true strain to a single inferred strain with similar abundance. However, some true strains had very low (<10%) relative abundance in all samples to which they were assigned in the simulation, and we expected reduced performance for these low abundance strains. Lower Jaccard similarity thresholds led to substantial false positives, with essentially all true strains having ≥3 matching inferred strains at thresholds ≤0.5 (**Figure 2A**). We defined >0.9 Jaccard similarity as a genotypic match between true and inferred strains to ensure high genotypic similarity while minimizing false positives. For strains that achieved a cumulative abundance over 1, almost all strains had a match at this cutoff (**Figure 2B**). At this threshold, 55% of simulated strains with cumulative abundance >0.1 had one or two matching inferred strains (**Figure 2A**, **Table S2**). This observation of many true strains having two inferred strains with close genotypic matches prompted us to investigate this phenomenon further.

**Figure 2.**
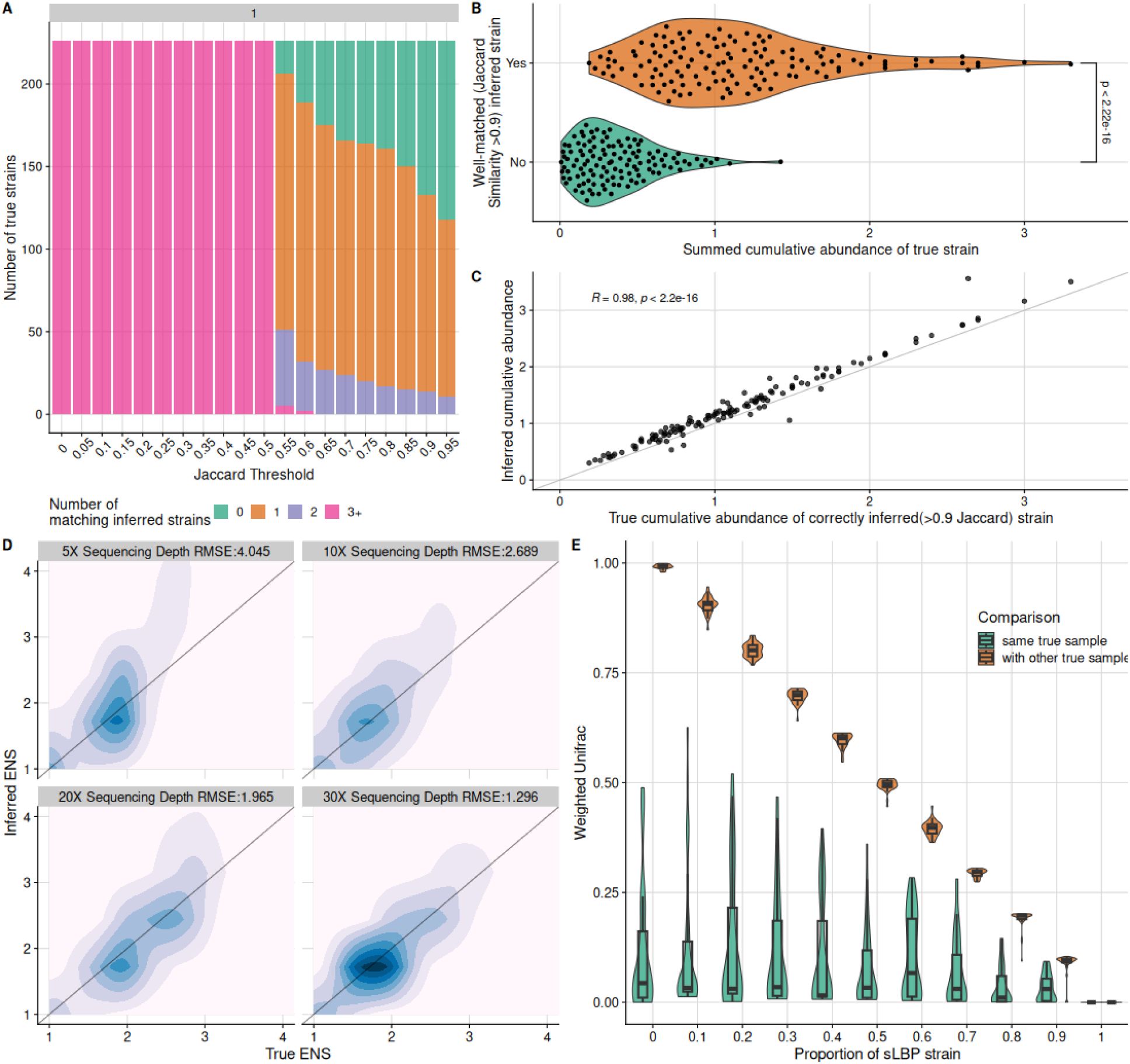
StrainFacts infers the genotype and composition of endogenous strains. A) Number of true strains that have the indicated numbers of matching inferred strains from the cohort at various Jaccard similarity thresholds. Data for the figure is shown for a single simulated cohort. B) Distribution of cumulative summed fractional abundances of true strains with genotypically well-matching (>0.9 Jaccard similarity) inferred strains compared to true strains that lacked well-matching inferred strains. A two sample t-test was conducted to compare the cumulative abundance means. Results are shown for an example simulated cohort. The inferred sLBP strain is excluded from this figure and analysis because of its much higher cumulative abundance due to presence in many samples. C) Correlation of true cumulative abundance for true strains with well-matching inferred strains (as in 2B, x-axis) and the inferred cumulative abundance of the matching inferred strains (y-axis). The inferred sLBP strain is not shown. D) Comparison of inferred effective number of strains (ENS) in each sample to the true ENS across sequencing depths. Root mean square error (RMSE) values are reported for the comparison of the effective number of strains at each depth. E) Comparison of weighted UniFrac dissimilarity between each sample’s inferred strain composition and its true strain composition (green) versus its dissimilarity with true strain compositions of other samples (green), visualized separately for each increment of sLBP proportion within the sample. Data are shown for an example simulation cohort.

In cases where a true strain had two closely matching inferred strains, we found that on average 80.33% of the inferred strain pairs were nearly identical (Jaccard similarity >0.99, **Table S3**). This indicated that StrainFacts had split metagenomic data from the single true strain into two nearly identical genotypes. In these cases, we collapsed the two near-identical genotypes into one by retaining the more abundant inferred strain and summing the abundances of both. The collapsed strains were used for all subsequent analysis.

We next examined why some true strains lacked an inferred strain with a close inferred match. True strains without a close match but with ≥ 10% abundance in at least one sample tended to have lower cumulative abundance (i.e., low summed abundance across all samplest median 0.316; IQR 0.169 - 0.504) compared to samples with close inferred matches (median 1.06; IQR 0.718 - 1.45), indicating that StrainFacts identified high-abundance strains well but was less effective for low-abundance strains (**Figure 2B**). Comparing the true and inferred summed abundances of matching paired and inferred strains showed a very strong correlation (*R* = 0.98 for all simulated cohorts), indicating excellent ability to determine strain abundance (**Figure 2C**). We then assessed how well strain alpha diversity was captured using the “effective number of strains” (ENS; e^(shannon)^) (Jost 2006). For each sample, we calculated true and inferred ENS, and evaluated their agreement and dependence on sample sequencing depth by calculating the RMSE at each depth (**Figure 2D**). At higher depths, StrainFacts accurately estimated richness with a lower RMSE. Performance declined at lower simulated sequencing depth, primarily due to overestimation of richness in samples with high true ENS at low sequencing depth.

To summarise the overall performance in inferring both strain genotype and abundance, we calculated the weighted UniFrac dissimilarity of the inferred and true strain composition and compared it to true strain compositions of other samples from the cohort (analyzed separately for each increment of simulated LBP abundance, **Figure 2E**). Weighted UniFrac is an ecological β diversity metric incorporating both relatedness (based on phylogenetic branch length distance – in our case calculated from Jaccard distance) and proportional abundance of members within two communities. The analysis revealed that dissimilarity between a sample’s inferred and true composition was consistently much lower than dissimilarity with other samples (**Figure 2E**). Samples with a high LBP abundance had lower weighted UniFrac dissimilarity overall because the LBP was well-inferred across all samples, but dissimilarity of each sample’s inferred strain composition with its true strain composition remained lower than dissimilarity with other samples (**Figure 2E**). The only exception was for samples with 100% sLBP abundance, which was anticipated because those samples are indistinguishable by this metric.

### StrainFacts identifies and quantifies a *Lactobacillus crispatus* LBP strain in spiked in to human vaginal samples

To assess StrainFacts’s ability to detect an LBP strain in clinical samples, we examined its performance in human vaginal microbiome samples with spike-ins of the *L. crispatus* strain CTV-05. CTV-05 is the bacterial component of the vaginally administered single-strain LBP LACTIN-V, which was shown in a randomized, placebo-controlled trial to prevent recurrent bacterial vaginosis (BV) following standard antibiotic treatment (Cohen 2020). We collected vaginal swab samples from ten participants with varying microbiota composition and performed nucleic acid extractions. Genomic DNA from a pure culture of CTV-05 was then added to each vaginal sample’s extracted DNA at two spike-in ratios (sample DNA:CTV-05 DNA ratios of 1,000:1 and 10:1). A control aliquot with no spike-in was retained for each sample, and all aliquots underwent short-read shotgun metagenomic sequencing (**Figure 3A**). Using VIRGO2 (France 2025), we quantified the relative abundances of the bacterial species in the metagenomic reads and observed a range of microbial community types including *L. crispatus*-dominant, *Lactobacillus iners*-dominant, and non-*Lactobacillus*-dominant - typical of the human vaginal microbiota spectrum **(Figure 3B**) (France 2020, Bloom 2022, Zhu 2024).

**Figure 3.**
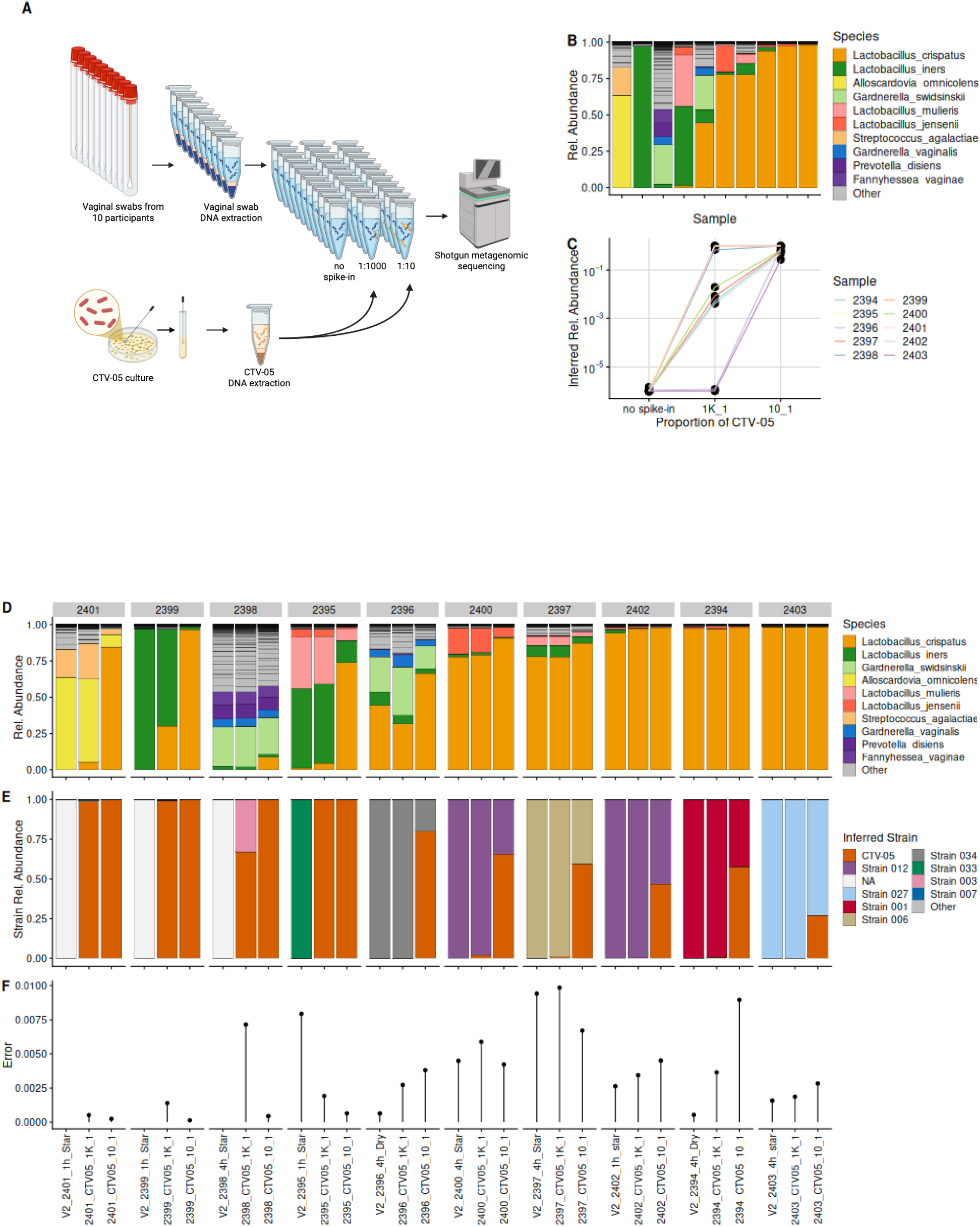
StrainFacts recovers spike-in LBP strain from human microbiota samples. A) Schematic of the protocol to spike LACTIN-V CTV-05 DNA into vaginal swabs. B) Microbiota composition of human vaginal swab samples used for CTV-05 spike-in experiments. Composition was determined from shotgun metagenomic sequences analyzed using VIRGO2 (France 2025). C) StrainFacts-inferred proportional abundance of the CTV-05 strain as a fraction of total *L. crispatus* within each sample. Aliquots of each sample were sequenced without spike-in or sequenced after spike-in of CTV-05 genomic DNA at the indicated ratio of sample DNA to genomic DNA. D) Relative abundances of bacterial species in each sample with or without CTV-05 spike-in at the indicated ratio. Abundances were determined from metagenomic sequences by VIRGO2. Data for the no-spike-in samples are the same as in panel B and are included here for comparison. E) StrainFacts-inferred proportional abundances of *Lactobacillus crispatus* strains as a fraction of total *L. crispatus* in each sample from C. F) Metagenotype errors reported by StrainFacts for strain estimates of the samples in E.

We generated a metagenotype from the sample reads using a database of core *L. crispatus* SNVs (**Figure S1**), and ran StrainFacts to infer *L. crispatus* strain abundances. Samples without sufficient *L. crispatus* coverage were excluded. Among the inferred strain genotypes, StrainFacts identified a strain with near-identical genotype as that of CTV-05. As expected, CTV-05 proportional strain abundance increased at higher spike-in concentrations (**Figure 3C**). In samples with low *L. crispatus* relative abundance, total *L. crispatus* relative abundance rose with increased spike-in concentration (**Figure 3D**). As expected, the inferred fractional abundance of CTV-05 increased within samples at higher spike-in concentration (**Figure 3E**). In samples where *L. crispatus* was rare or absent, StrainFacts inferred the strain composition to consist entirely of the spiked in LBP strain. However, in sample 2398, another *L. crispatus* strain (Strain 003) was inferred (**Figure 3E**). This sample had a higher metagenotype error (**Figure 3F**), which is a reflection of how well the StrainFacts inference matched the true metagenotypee, suggesting that Strain 003 was spurious. In samples with higher endogenous *L. crispatus*, endogenous *L. crispatus* strains were inferred along with spiked-in CTV-05, but proportions of CTV-05 rose in the samples at greater concentrations of spike-in (Figure3E).

Within each parent sample that had endogenous *L. crispatus*, StrainFacts consistently identified the same endogenous strain across the parent and respective spiked-in samples. Endogenous strains were unique to each parent sample, with the exception of samples 2400 and 2402, where Strain 12 was inferred in both (**Figure 3E**). This could either reflect an inference error or the true presence of highly similar endogenous strains in both samples. To explore why some samples had higher metagenotype errors, we calculated the spread of the sequencing depths across all database positions for each sample. The results demonstrated that samples with lower sequencing depths often had higher metagenotype errors (**Figure S2**).

## Discussion

Live biotherapeutic products (LBPs) promote health by introducing one or more microbial strains into a community that may also contain endogenous strains of the same species, yet full characterization of LBP effects requires validated methods to distinguish and quantify both administered and endogenous strains. Here, we validated the ability of StrainFacts to track and quantify an LBP strain in both simulated microbiota samples and human vaginal metagenome samples spiked with the *L. crispatus* strain from the vaginal LBP LACTIN-V (Smith 2022, Cohen 2020). StrainFacts was selected for this analysis because of its potential to identify, distinguish, and quantify both LBP and endogenous strains of a species in the same samples. Ideally, a strain estimation tool would accurately infer both genotype and abundance of a strain matching the true genotype of the LBP strain as well as the genotypes and abundances of any endogenous strains, including determining whether distinct samples contain identical strains (e.g., the same LBP strain in multiple recipients or the same endogenous strain in serial samples from a single participant). Our findings indicate that StrainFacts effectively fulfills these functions, demonstrating its utility for characterizing strain-level dynamics in LBP trials.

In addition to demonstrating the applicability of a specific tool for tracing strains in clinical samples, our study also illustrates the value of real or simulated LBP trial data for validating strain-tracing tool performance in general. Strain-tracing in metagenomic samples from observational clinical cohorts or other sources without defined live microbial treatment typically involves profiling communities presumed to contain many endogenous strains whose true genotypes and abundances are unknown. Since LBP treatment introduces one or more strains with known (or knowable) genotypes into otherwise undefined communities, comparing verifiable LBP strain genotype and abundance to a strain-tracing tool’s output can provide a valuable “ground truth” for assessing tool accuracy.

Further studies should explore whether endogenous strains identified by StrainFacts and their relative abundances match true bacterial isolates cultured from samples. Although StrainFacts performed well with real-word human vaginal samples, we observed one instance where the same endogenous *L. crispatus* strain was inferred in two different participants despite low metagenotype error. This may reflect either an error by StrainFacts or that these two participants may indeed have been colonized by an identical *L. crispatus* strain or highly similar strains.

While we found this approach to be effective in quantifying a single-strain LBP, our study does have limitations and further method development and validation is needed. Accurate strain-tracing for LBP formulations containing multiple strains of a single species or strains from multiple species will be more complex and require further validation. In addition, our findings may not fully extend to LBPs or microbial treatments targeting microbiota r other than the those in the human vagina, which is known to have overall less diversity than many other human and non-human microbial communities. We found that high sequencing depth of the target species was required, and even at high depths at least ∼10% of the LBP strain was needed within the target species for confident detection. While this threshold is reasonable for assessing LBPs that achieve robust colonization, it may limit applicability for products that achieve lower colonization or for tracking subtle shifts inf endogenous strains over time. In addition, access to numerous high-quality reference genomes of the species of interest is required for generating the GT-Pro database that StrainFacts utilizes. This requirement could limit the use of StrainFacts for LBPs or other defined live microbial treatments containing species with few sequenced representatives, although species used in such products are likely to be well-characterized and well-represented in genome databases.

## Methods

### A simulation of metagenomic samples to validate StrainFacts inferences

To validate StrainFacts for use in estimating the presence, abundance, and genotypes of endogenous bacterial strains and of an administered LBP strain within metagenomic samples, we investigated its ability to infer strains in a simulated, single-arm LBP treatment cohort. We first used StrainFacts to generate a metagenotype of 100 simulated participants, each with 4 distinct simulated samples. The resulting count matrix was then adjusted for each participant to be assigned 1 to 4 simulated endogenous strains from a single species, with each strain possessing a unique simulated genotype consisting of a pattern of SNVs varying across 10,000 biallelic sites. The four simulated samples for each participant were assigned to contain that participant’s 1-4 endogenous strains at abundances which varied from sample to sample. No endogenous strains were simulated to be shared between participants. Samples were simulated to have sequencing depth (i.e., SNV site coverage) of 5x, 10x, 20x, or 30x before the introduction of the LBP strain. To simulate LBP treatment, a single simulated LBP (sLBP) strain genotype was proportionally “spiked in’’ to each sample at a per-sample LBP strain abundance stochastically varying from 0% to 100% (in 10% increments) by adjusting the counts at each SNV site within the metagenotype. Thus, each simulated sample consisted of a defined mixture of 1-4 participant-specific endogenous strains and one shared sLBP strain, with each strain in each sample having a known (“true”) genotype and abundance. Employing these adjusted metagenotypes as input, we then used StrainFacts to generate inferred strain genotypes and abundances for each sample. We performed this process 10 times, producing 10 independent simulated cohorts, to investigate consistency and accuracy of the estimations. We used the same sLBP strain genotype for all 10 simulated cohorts, but allowed endogenous strain genotypes to vary between simulations. (**Figure 1A**).

### Sample Collection

Human vaginal swab samples were collected from 10 premenopausal participants enrolled in the V2 cohort, which is an IRB-approved cohort at Massachusetts General Hospital (MGH IRB Protocol 2014P001066) that enrolls women presenting for gynecological care at MGH who provided written informed consent for sample collection (Mitchell 2020). Women with HIV or those who are otherwise immunocompromised are not enrolled and participants received no monetary compensation. The swabs were stored at -80C in StarSwab media prior to processing.

### Nucleic Acid Extraction from human vaginal swabs and culture CTV-05

Total nucleic acids (TNA, including gDNA) from swabs were extracted using a phenol-chloroform method, which includes a bead beating process to disrupt bacteria (Anahtar et. al. 2016), with a modification for 96-well plate (Hemmerling et al. 2025). In short, an aliquot of 200uL of sample solution was transferred into cluster tubes containing ∼200mg sterile 0.1mm glass beads, 200uL of phenol chloroform, 80uL of 20% sodium dodecyl sulphate and 8.14uL of EDTA. After incubating at 4C for 5 min, samples were homogenized on a bead beater and centrifuged at 2800xg for 15 min at 4C. The top aqueous phase was transferred to 96-well deep plates, and 300uL of -20C isopropanol and 30uL of sodium acetate were added to precipitate TNA overnight at -20C. The plate was centrifuged at 2800xg for 30 minutes at 4C and supernatant was removed, and the TNA pellet was washed with 300uL of 100% ethanol. After centrifuge at 2800xg for 15 minutes at 4C, ethanol was removed and the plate left open to dry at room temperature. The TNA pellet was resuspended in 85uL of TE buffer and stored at -80C. TNA from a pure culture of the Lactin-V strain CTV-05 was extracted with the same protocols as the swabs, except that the CTV-05 pellet was suspended in 200uL of Tris-EDTA buffer instead of StarSwab media.

### Spiking in Lactin-V CTV-05 DNA to swab samples

DNA from each sample and from CTV-05 was quantified with SYBR Green I. The DNA sample of Lactin-V CTV-05 was spiked into swab DNA samples concentration ratios of 1:10 (100_1) and 1:1000 (1K_1), respectively by adding CTV-05 DNA to swab DNA samples before library preparation.

### Shotgun Metagenomic Library Preparation and Sequencing

Shotgun library was prepared following a modified protocol of Baym et. al (2015), using the Nextera DNA Library Preparation Kit (Illumina) and KAPA HiFi Library Amplification Kit (Kapa Biosystems). In brief, gDNA from each sample was standardized to concentration of 1ng/uL after quantification with SYBR Green I, followed by simultaneous fragmentation and sequencing adaptor incorporation by mixing 1ng gDNA (1uL) with 1.25uL TD buffer and 0.25uL TDE1 provided in the Nextera kit and incubating for 9 minutes at 55°C. Tagmented DNA fragments were amplified in PCR using the KAPA high fidelity library amplification reagents, with Illumina adaptor sequences and sample barcodes incorporated in primers. PCR products were pooled, purified with magnetic beads and paired-end sequenced on Illumina NovaSeq X with a 300-cycle kit.

### Shotgun Metagenomic Read Processing and Taxonomic Profiling

Cutadapt v4.6 was used to remove adapters from raw reads. Quality filtering was performed with a quality threshold of Q20 and minimum read length of 50 bp using Sickle-trim v1.33. Human reads were removed using BBDuk from the BBtools suite Human-filtered metagenomic reads from both the parent and spiked-in samples were processed through the VIRGO2 pipeline to quantify relative abundances of bacterial species (France 2025).

### Construction of GT-pro *L. crispatus* database

137 *L. crispatus* genomes of high quality were used to construct a database of core biallelic SNPs. MAAST was used to identify the core biallelic SNPs, where core was defined as present in 95% of genomes (131/137), and SNP was defined as a minimum allele frequency of 2% (2/137). The MAAST SNPs were then built into a database using the GTPro build command.

### Strain inference with MAAST, GT-Pro, and StrainFacts

Strain-level inference was performed using MAAST (Shi 2023), GT-Pro (Shi 2022), and StrainFacts in a Snakemake pipeline. StrainFacts requires a user-defined parameter for the expected number of strains in the total sampleset (which we set at 300 strains for our simulated cohort analysis), so StrainFacts produced 300 inferred strain genotypes and inferred an abundance for each of the 300 in each sample, but we anticipated strains should be inferred at near-zero (e.g., <10^−5^) abundance for samples in which they were not truly present.

## Acknowledgments

We thank Laura Symul and Michael France for helpful discussions. Funding was obtained by

D.S.K. from the Gates Foundation (INV-033690). S.M.B. was partially supported by NIH grant 1K08AI171166.

## Author Contributions

J.S. and J.E. conceptualized the research with input from S.M.B. and D.S.K.; S.M.B., J.X., and C.M.M. generated samples and performed experiments; J.S. and J.E. performed data analysis with input from S.M.B.; J.S. and J.E. led writing of the manuscript with contributions from S.M.B., J.X, and D.S.K.; All authors provided critical feedback on methods, results, analysis, and writing.

## Data and code availability

Code for simulations are available at https://github.com/jshih08/sfacts_lbp_simulation. Shotgun metagenomic reads are available at BioProject PRJNA1306570.

## Supplemental Figures

**Figure S1.**
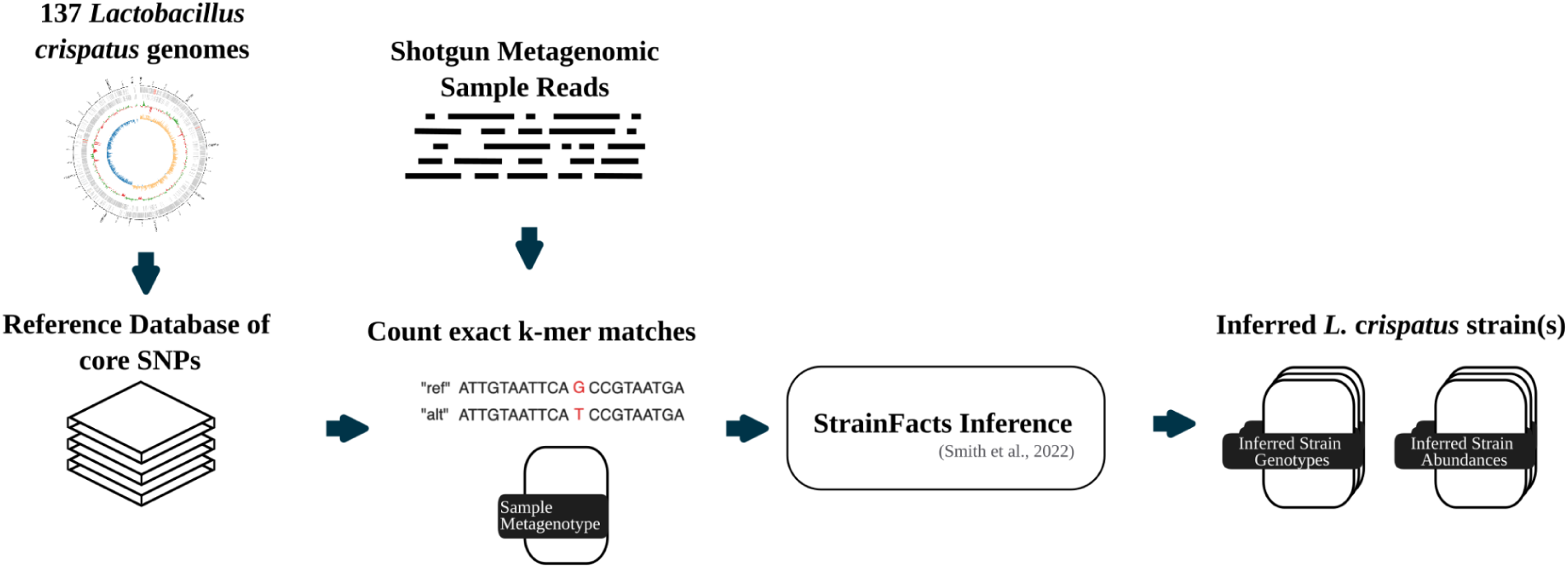
*Lactobacillus crispatus* LBP strain quantification using a StrainFacts workflow.

**Figure S2.**
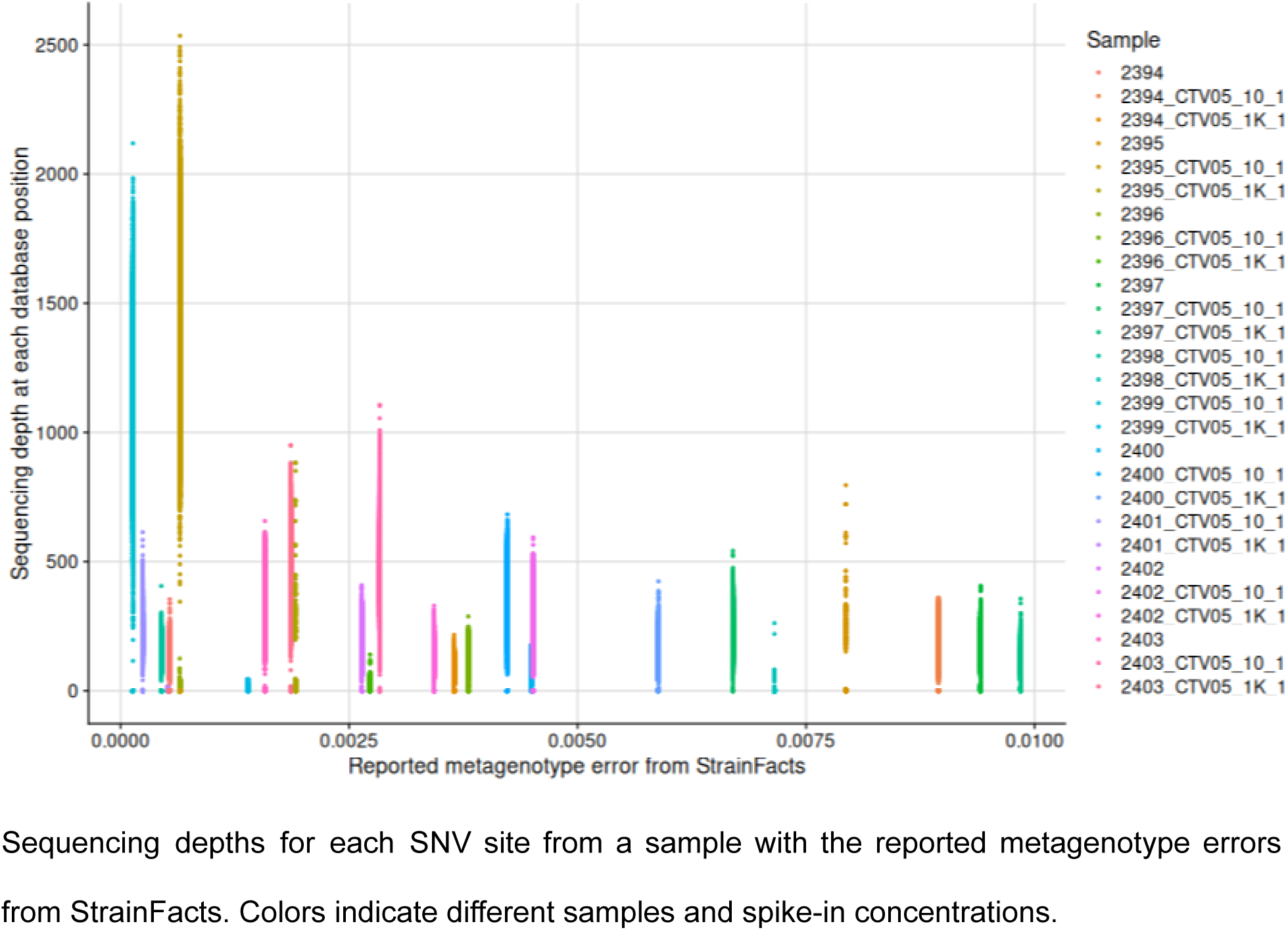
Sequencing depths are associated with metagenotype error.

---

**Table S1.**
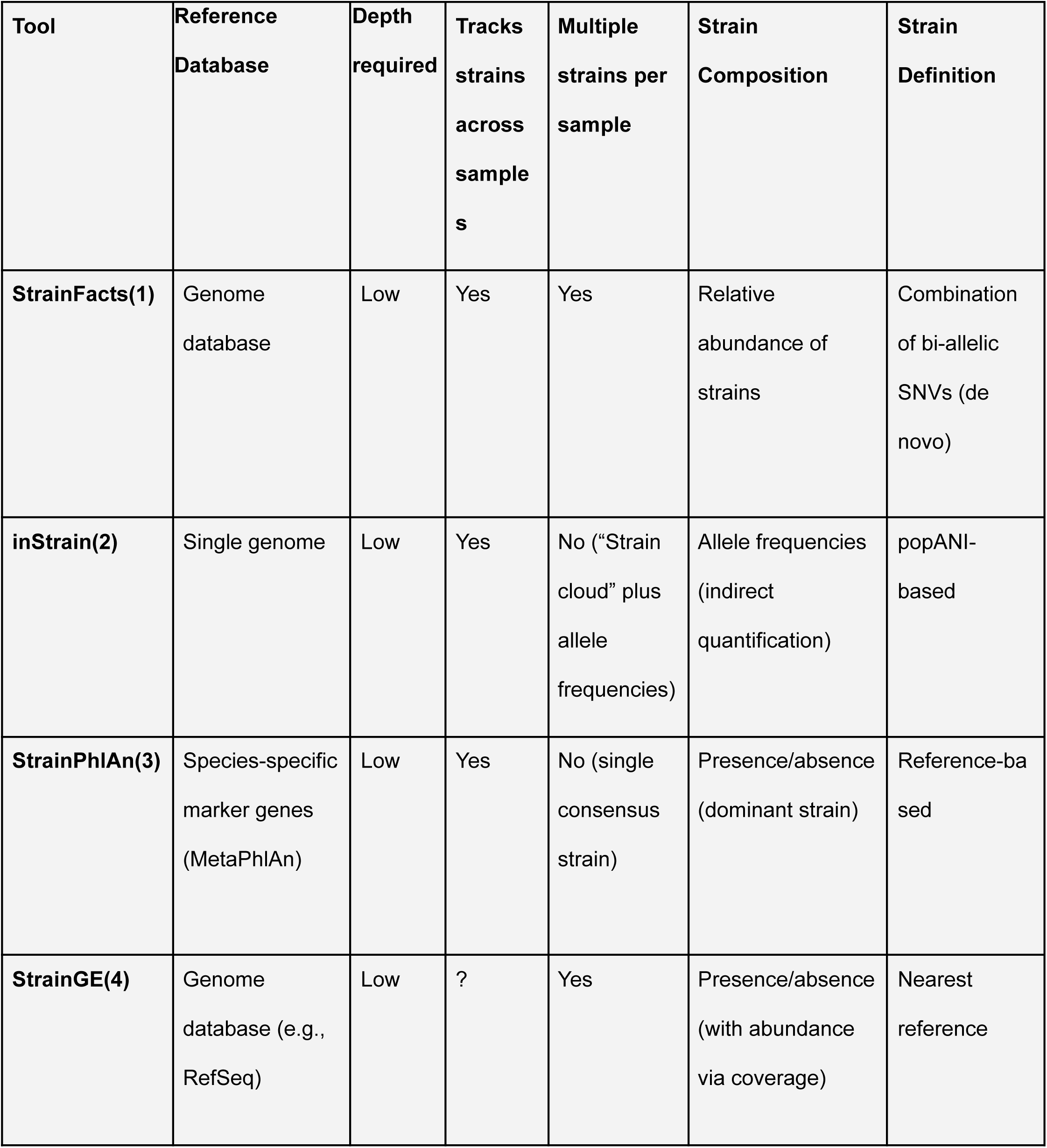
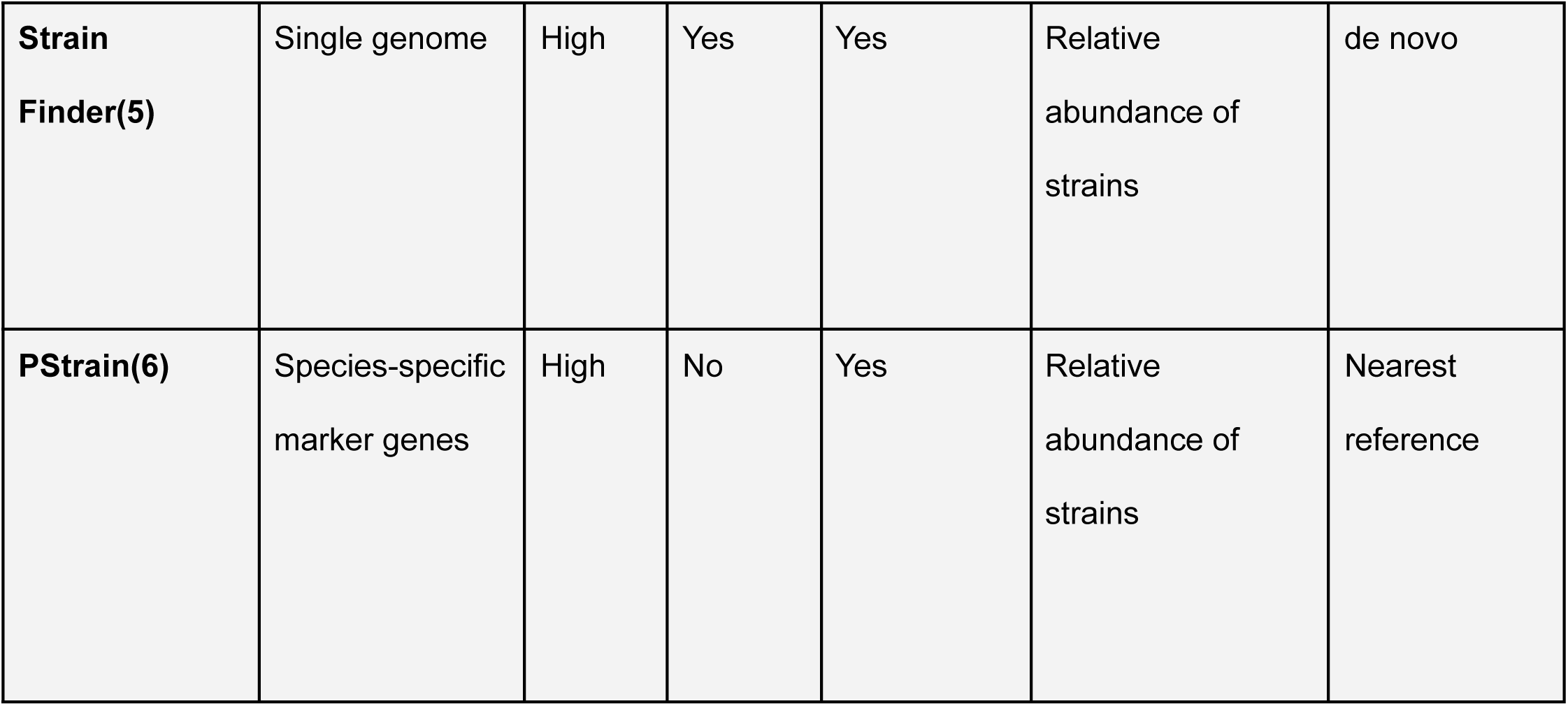
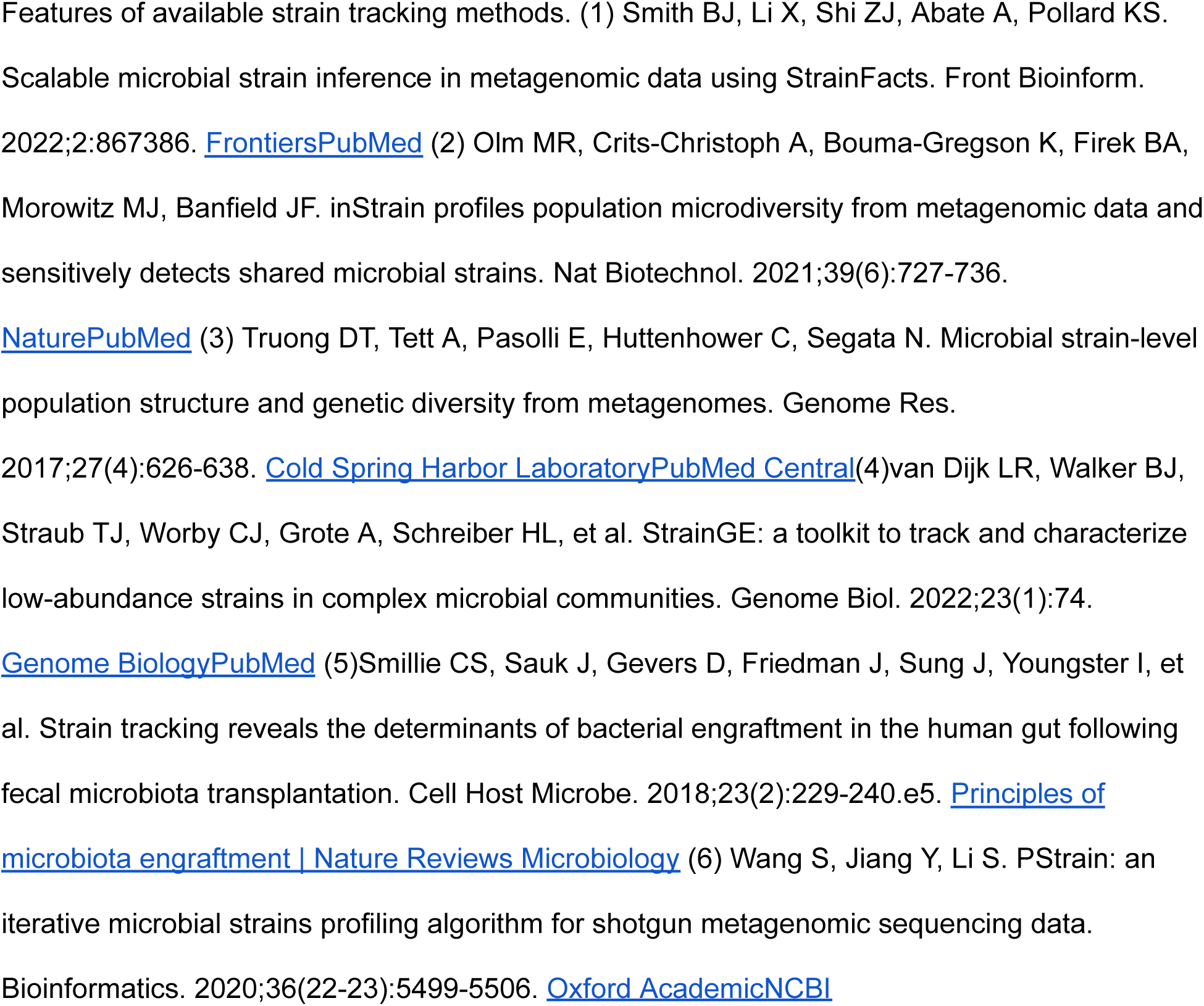
comparison of StrainFacts with other strain inference methods.

**Table S2.** Number of inferred strains of each true strain across simulated cohorts.

|  | **Cohort**<br>**1, N =**<br>**226<sup>1</sup>** | **Cohort**<br>**2, N =**<br>**219<sup>1</sup>** | **Cohort**<br>**3, N =**<br>**223<sup>1</sup>** | **Cohort**<br>**4, N =**<br>**225<sup>1</sup>** | **Cohort**<br>**5, N =**<br>**226<sup>1</sup>** | **Cohort**<br>**6, N =**<br>**229<sup>1</sup>** | **Cohort**<br>**7, N =**<br>**218<sup>1</sup>** | **Cohort**<br>**8, N =**<br>**228<sup>1</sup>** | **Cohort**<br>**9, N =**<br>**227<sup>1</sup>** | **Cohort**<br>**10, N =**<br>**218<sup>1</sup>** |
| --- | --- | --- | --- | --- | --- | --- | --- | --- | --- | --- |
| 0 | 93<br>(41%) | 104<br>(47%) | 100<br>(45%) | 95<br>(42%) | 92<br>(41%) | 101<br>(44%) | 96<br>(44%) | 110<br>(48%) | 108<br>(48%) | 106<br>(49%) |
| 1 | 119<br>(53%) | 108<br>(49%) | 115<br>(52%) | 121<br>(54%) | 126<br>(56%) | 120<br>(52%) | 108<br>(50%) | 107<br>(47%) | 112<br>(49%) | 103<br>(47%) |
| 2 | 14<br>(6.2%) | 7<br>(3.2%) | 8<br>(3.6%) | 9<br>(4.0%) | 8<br>(3.5%) | 8<br>(3.5%) | 14<br>(6.4%) | 11<br>(4.8%) | 7<br>(3.1%) | 9<br>(4.1%) |
| <sup>1</sup> n (%) |  |  |  |  |  |  |  |  |  |  |

Number (percent) of inferred strains that genotypically match each true strain with Jaccard similarity >0.9 for each of the 10 simulated cohorts (number of strains analyzed per cohort is indicated in the header of each column). Analysis is restricted to true strains with ≥10% abundance in at least one sample.

**Table S3.** Percentage of well-matched inferred strains that are near identical.

| Simulated Cohort | % of true strains with well-matching pairs of inferred that are near identical (Jaccard Similarity > 0.99) |
| --- | --- |
| 1 | 71.43% |
| 2 | 92.86% |
| 3 | 81.25% |
| 4 | 88.89% |
| 5 | 75.00% |
| 6 | 81.25% |
| 7 | 67.86% |
| 8 | 77.27% |
| 9 | 78.57% |
| 10 | 88.89% |

Percentage of true strains with two well-matching inferred strains (>0.9 Jaccard similarities of each inferred strain to the true strain) in which the pair of inferred strains were nearly identical to each other (>0.99 Jaccard similarity between the inferred strains).

## References

Bloom, S. M., Mafunda, N. A., Woolston, B. M., Hayward, M. R., Frempong, J. F., Abai, A. B., Xu, J., Mitchell, A. J., Westergaard, X., Hussain, F. A., Xulu, N., Dong, M., Dong, K. L., Gumbi, T., Ceasar, F. X., Rice, J. K., Choksi, N., Ismail, N., Ndung’u, T., … Kwon, D. S. (2022). Cysteine dependence of Lactobacillus iners is a potential therapeutic target for vaginal microbiota modulation. Nature Microbiology, 7(3), 434–450. 10.1038/s41564-022-01070-7

Cohen, C. R., Wierzbicki, M. R., French, A. L., Morris, S., Newmann, S., Reno, H., Green, L., Miller, S., Powell, J., Parks, T., & Hemmerling, A. (2020). Randomized Trial of Lactin-V to Prevent Recurrence of Bacterial Vaginosis. New England Journal of Medicine, 382(20), 1906–1915. 10.1056/NEJMoa1915254

Du Toit, A. (2018). Principles of microbiota engraftment. Nature Reviews Microbiology, 16(4), 186–186. 10.1038/nrmicro.2018.29

France, M. T., Chaudry, I., Rutt, L., Quain, M., Shirtliff, B., McComb, E., Maros, A., Alizadeh, M., Hussain, F. A., Elovitz, M. A., Relman, D. A., Rahman, A., Brotman, R. M., Price, J., Kassaro, M., Holm, J. B., Ma, B., & Ravel, J. (2025). VIRGO2: Unveiling the Functional and Ecological Complexity of the Vaginal Microbiome with an Enhanced Non-Redundant Gene Catalog. 10.1101/2025.03.04.641479

France, M. T., Ma, B., Gajer, P., Brown, S., Humphrys, M. S., Holm, J. B., Waetjen, L. E., Brotman, R. M., & Ravel, J. (2020). VALENCIA: a nearest centroid classification method for vaginal microbial communities based on composition. Microbiome, 8(1), 166. 10.1186/s40168-020-00934-6

Hansen, R., Sanderson, I. R., Muhammed, R., Allen, S., Tzivinikos, C., Henderson, P., Gervais, L., Jeffery, I. B., Mullins, D. P., O’Herlihy, E. A., Weinberg, J. D., Kitson, G., Russell, R. K., & Wilson, D. C. (2021). A Double-Blind, Placebo-Controlled Trial to Assess Safety and Tolerability of (Thetanix) Bacteroides thetaiotaomicron in Adolescent Crohn’s Disease. Clinical and Translational Gastroenterology, 12(1), e00287. 10.14309/ctg.0000000000000287

Jost, L. (2006). Entropy and diversity. Oikos, 113(2), 363–375. 10.1111/j.2006.0030-1299.14714.x

Khanna, S., Pardi, D. S., Jones, C., Shannon, W. D., Gonzalez, C., & Blount, K. (2021). RBX7455, a Non-frozen, Orally Administered Investigational Live Biotherapeutic, Is Safe, Effective, and Shifts Patients’ Microbiomes in a Phase 1 Study for Recurrent *Clostridioides difficile* Infections. Clinical Infectious Diseases, 73(7), e1613–e1620. 10.1093/cid/ciaa1430

Mitchell, C. M., Watson, L., Mitchell, A. J., Hyrien, O., Bergerat, A., Valint, D. J., Pascale, A., Hoffman, N., Srinivasan, S., & Fredricks, D. N. (2020). Vaginal Microbiota and Mucosal Immune Markers in Women With Vulvovaginal Discomfort. Sexually Transmitted Diseases, 47(4), 269–274. 10.1097/OLQ.0000000000001143

Olm, M. R., Crits-Christoph, A., Bouma-Gregson, K., Firek, B. A., Morowitz, M. J., & Banfield, J. F. (2021a). inStrain profiles population microdiversity from metagenomic data and sensitively detects shared microbial strains. Nature Biotechnology, 39(6), 727–736. 10.1038/s41587-020-00797-0

Olm, M. R., Crits-Christoph, A., Bouma-Gregson, K., Firek, B. A., Morowitz, M. J., & Banfield, J. F. (2021b). inStrain profiles population microdiversity from metagenomic data and sensitively detects shared microbial strains. Nature Biotechnology, 39(6), 727–736. 10.1038/s41587-020-00797-0

Shi, Z. J., Dimitrov, B., Zhao, C., Nayfach, S., & Pollard, K. S. (2022). Fast and accurate metagenotyping of the human gut microbiome with GT-Pro. Nature Biotechnology, 40(4), 507–516. 10.1038/s41587-021-01102-3

Shi, Z. J., Nayfach, S., & Pollard, K. S. (2023). Maast: genotyping thousands of microbial strains efficiently. Genome Biology, 24(1), 186. 10.1186/s13059-023-03030-8

Smillie, C. S., Sauk, J., Gevers, D., Friedman, J., Sung, J., Youngster, I., Hohmann, E. L., Staley, C., Khoruts, A., Sadowsky, M. J., Allegretti, J. R., Smith, M. B., Xavier, R. J., & Alm, E. J. (2018). Strain Tracking Reveals the Determinants of Bacterial Engraftment in the Human Gut Following Fecal Microbiota Transplantation. Cell Host & Microbe, 23(2), 229–240.e5. 10.1016/j.chom.2018.01.003

Smith, B. J., Li, X., Shi, Z. J., Abate, A., & Pollard, K. S. (2022b). Scalable Microbial Strain Inference in Metagenomic Data Using StrainFacts. Frontiers in Bioinformatics, 2, 867386. 10.3389/fbinf.2022.867386

Truong, D. T., Tett, A., Pasolli, E., Huttenhower, C., & Segata, N. (2017a). Microbial strain-level population structure and genetic diversity from metagenomes. Genome Research, 27(4), 626–638. 10.1101/gr.216242.116

Truong, D. T., Tett, A., Pasolli, E., Huttenhower, C., & Segata, N. (2017b). Microbial strain-level population structure and genetic diversity from metagenomes. Genome Research, 27(4), 626–638. 10.1101/gr.216242.116

Truong, D. T., Tett, A., Pasolli, E., Huttenhower, C., & Segata, N. (2017c). Microbial strain-level population structure and genetic diversity from metagenomes. Genome Research, 27(4), 626–638. 10.1101/gr.216242.116

van Dijk, L. R., Walker, B. J., Straub, T. J., Worby, C. J., Grote, A., Schreiber, H. L., Anyansi, C., Pickering, A. J., Hultgren, S. J., Manson, A. L., Abeel, T., & Earl, A. M. (2022a). StrainGE: a toolkit to track and characterize low-abundance strains in complex microbial communities. Genome Biology, 23(1), 1–27. 10.1186/s13059-022-02630-0

van Dijk, L. R., Walker, B. J., Straub, T. J., Worby, C. J., Grote, A., Schreiber, H. L., Anyansi, C., Pickering, A. J., Hultgren, S. J., Manson, A. L., Abeel, T., & Earl, A. M. (2022b). StrainGE: a toolkit to track and characterize low-abundance strains in complex microbial communities. Genome Biology, 23(1), 1–27. 10.1186/s13059-022-02630-0

Vermeire, S., Dewint, P., Vansteelant, M., Peterka, M., Štěpek, D., Kierkuś, J., Wiernicka, A., Napora, P., Wolański, Ł., Kopoń, A., Magro, F., Pinheiro, I., Possemiers, S., Haazen, L., & Bolca, S. (2024). P658 Safety and efficacy of MH002, an optimized live biotherapeutic product, for the treatment of mild to moderate ulcerative colitis: a first-in-disease, double-blind, randomized clinical trial. Journal of Crohn’s and Colitis, 18(Supplement_1), i1243–i1244. 10.1093/ecco-jcc/jjad212.0788

Wang, S., Jiang, Y., & Li, S. (2021a). PStrain: an iterative microbial strains profiling algorithm for shotgun metagenomic sequencing data. Bioinformatics, 36(22–23), 5499–5506. 10.1093/bioinformatics/btaa1056

Wang, S., Jiang, Y., & Li, S. (2021b). PStrain: an iterative microbial strains profiling algorithm for shotgun metagenomic sequencing data. Bioinformatics, 36(22–23), 5499–5506. 10.1093/bioinformatics/btaa1056

Zhu, M., Frank, M. W., Radka, C. D., Jeanfavre, S., Xu, J., Tse, M. W., Pacheco, J. A., Kim, J. S., Pierce, K., Deik, A., Hussain, F. A., Elsherbini, J., Hussain, S., Xulu, N., Khan, N., Pillay, V., Mitchell, C. M., Dong, K. L., Ndung’u, T., … Kwon, D. S. (2024). Vaginal Lactobacillus fatty acid response mechanisms reveal a metabolite-targeted strategy for bacterial vaginosis treatment. Cell, 187(19), 5413–5430.e29. 10.1016/j.cell.2024.07.029

